# Integrated Post-Translational Modification (PTM) Proteomics Reveals Early Redox and Mitochondrial Remodeling Following Ref-1 Inhibition in Pancreatic Cancer

**DOI:** 10.64898/2026.08.15.745014

**Authors:** Silpa Gampala, Xiaolu Li, Jesse B Trejo, Marina A Gritsenko, Rosalie K Chu, Wei-Jun Qian, Elizabeth Sierra Potchanant, Melissa L. Fishel, Tong Zhang, Mark R. Kelley

## Abstract

**Background:** Apurinic/apyrimidinic endonuclease 1/redox factor-1 (Ref-1/APE1) is a central regulator of redox-dependent transcriptional signaling that promotes pancreatic ductal adenocarcinoma (PDAC) progression, therapeutic resistance, and metabolic adaptation. While pharmacologic inhibition of Ref-1 suppresses tumor growth and alters cellular metabolism, immediate molecular events linking Ref-1 inhibition to downstream cellular adaptation remain poorly understood. We therefore sought to characterize proteome-wide signaling responses induced by second-generation Ref-1 redox inhibitor, APX2014.

**Methods:** We applied an integrated multiplexed proteomics workflow to simultaneously quantify global protein abundance together with cysteine oxidation, phosphorylation, and lysine acetylation in Pa03C PDAC cells following acute treatment (30–120 min) with selective Ref-1 redox inhibitor APX2014. Differential post-translational modification (PTM) analysis, pathway enrichment, structural mapping of regulated sites, and functional mitochondrial substrate utilization assays were performed to define early signaling responses.

**Results:** APX2014 induced rapid and extensive remodeling of PTM landscape while producing minimal changes in global protein abundance. Cysteine oxidation represented the earliest and most sustained response, accompanied by widespread phosphorylation and delayed lysine acetylation. Integrated pathway analyses identified mitochondrial translation, respiratory electron transport, TCA cycle metabolism, and mitochondrial redox homeostasis as the earliest and most consistently regulated processes. Functional mitochondrial assays confirmed impaired utilization of TCA cycle substrates following APX2014 treatment. Coordinated PTM remodeling was observed on Ref-1-associated signaling proteins, including NF-κB1 and p53, revealing simultaneous regulation of oxidation, phosphorylation, and acetylation within functionally important domains. Early redox-sensitive protein networks were also associated with subsequent disruption of mitotic organization.

**Conclusions:** Integrated multi-PTM proteomics reveals that pharmacologic Ref-1 redox inhibition rapidly rewires regulatory signaling networks before detectable changes in protein abundance. Our findings identify mitochondrial redox remodeling as an early consequence of Ref-1 inhibition, providing systems-level insight into how Ref-1-targeted therapies disrupt metabolic and stress-adaptive programs in pancreatic cancer. This work establishes a framework for understanding the molecular basis of Ref-1-directed therapeutics and highlights integrated PTM profiling as a powerful strategy for defining early drug response mechanisms. These findings provide a strong translational rationale for advancing next-generation Ref-1 redox inhibitors such as APX2014, developed from the first-in-class inhibitor APX3330 currently in clinical trials, and underscore the broader therapeutic potential of targeting Ref-1 redox signaling in pancreatic cancer.

## Background

Pancreatic ductal adenocarcinoma (PDAC) remains one of the deadliest cancers, with few treatment options, often diagnosed late, and quick progression or inherent resistance to therapy. Although FOLFIRINOX, gemcitabine/nab-paclitaxel, and emerging KRAS inhibitors can extend treatment options for some patients, durable responses remain rare and therapeutic resistance commonly develops (1–3). A key factor in PDAC’s aggressiveness is its ability to adapt to cellular and environmental stress by regulating redox balance, mitochondrial function, DNA repair, and survival signals. These adaptive processes enable PDAC cells to survive in low oxygen, limited nutrients, oxidative stress, inflammation, and damage from treatments, allowing tumors to persist under conditions that would normally harm cells (4).

Post-translational modifications (PTMs) provide tumor cells fast and reversible regulation, reshaping signaling networks without major changes in protein levels (5–7). PTMs such as phosphorylation, acetylation, ubiquitination, SUMOylation, and cysteine oxidation control protein activity, stability, localization, and interactions. In PDAC, these modifications are connected to tumor growth, metabolic flexibility, stromal interactions, DNA repair, inflammatory signaling, and resistance to therapy (8). Importantly, PTMs signal within connected regulatory networks rather than as isolated events (9). Coordinated modification patterns across signaling proteins generate complex regulatory states that influence pathway activity, protein function, and cellular adaptation to stress or therapeutic intervention. This highlights the importance of multi-PTM profiling for understanding early adaptive signaling in PDAC.

Apurinic/apyrimidinic endonuclease 1/Redox factor-1 (APE1/Ref-1) is a multifunctional protein that combines DNA base excision repair with redox regulation of transcription factors. Through its redox activity, Ref-1 influences the DNA-binding and transcriptional activity of factors involved in cancer cell survival, inflammatory signaling, stress adaptation, and therapeutic resistance, including NF-κB, HIF-1α, STAT3, AP-1, and p53 (10). Because these pathways are key to PDAC progression, pharmacological inhibition of Ref-1 redox signaling has become a promising therapeutic approach (11). APX2014 is a small-molecule inhibitor designed to selectively block Ref-1–dependent redox signaling while maintaining its DNA repair endonuclease function (11). Previous studies have demonstrated that Ref-1 redox inhibition can suppress tumor-promoting transcriptional programs, hinder PDAC cell growth and survival, and impact pathways related to oxidative stress, metabolism, and therapeutic sensitivity (12–14). However, the early biochemical events linking Ref-1 redox inhibition to downstream PTM signaling networks in PDAC remain poorly understood.

This gap is especially important because Ref-1 redox signaling acts upstream of several transcriptional and signaling pathways that are themselves controlled by PTMs. Inhibiting Ref-1 redox activity could therefore lead to immediate biochemical effects that impact multiple pathways downstream of Ref-1, including changes in cysteine oxidation, kinase signaling, transcriptional activity, mitochondrial function, and metabolic adaptation. These early responses are likely to occur before detectable changes in overall transcription factor activity and may define the primary cellular response to APX2014. Capturing this immediate regulatory window is crucial to distinguish direct vs indirect effects of Ref-1 redox blocking from later secondary changes.

Mass spectrometry-based proteomics offers a powerful approach to understanding early adaptive responses across the entire proteome. Quantitative PTM proteomics enables site-specific measurement of regulatory events that are not detectable from protein abundance alone. Phosphoproteomics can identify kinase-driven signaling pathways (9, 15), acetylomics reveals changes in chromatin, metabolism, and mitochondrial proteins, and cysteine-focused redox proteomics directly measures redox-sensitive thiol oxidation events (16, 17). Combining these PTM events with overall protein abundance provides a systems-level view of how cancer cells quickly reorganize signaling and stress-response pathways in response to pharmacologic disturbances. Importantly, quantifying multiple PTM classes in parallel is essential because PTMs frequently co-occur and exhibit crosstalk on the same proteins. Yet most PTM workflows are performed separately, requiring additional sample input and introducing variability that complicates integrative interpretation. To address this, our group developed an integrative workflow that enables multiplexed quantification of global protein abundance together with phosphorylation, acetylation, and cysteine thiol oxidation from the same samples (18). This unified, automation-friendly approach improves throughput and consistency, enabling time-resolved, systems-level mapping of coordinated multi-PTM responses to drug perturbation.

Here, we employed our integrated multi-PTM proteomics approach to characterize the early cellular response to APX2014 in pancreatic cancer cell line, Pa03C. Using multiplexed TMT18 labeling combined with parallel enrichment workflows, global protein levels were measured along with site-specific cysteine oxidation, phosphorylation, and lysine acetylation over an acute treatment period (30 min – 2 h). This approach enabled us to differentiate rapid redox-related events from later adaptive responses and to identify signaling nodes with coordinated PTM changes. Our findings indicate that APX2014 quickly elevates cysteine oxidation and causes widespread phosphorylation, with relatively delayed changes in acetylation, while overall protein levels remain mostly stable during this early phase. Additionally, we observe coordinated PTM regulation of Ref-1–linked signaling proteins, such as NF-κB1 and p53, and highlight early mitochondrial remodeling as a key feature of the APX2014 response in PDAC cells.

## Methods

### Cell Culture and Treatment

Pancreatic cancer cell line Pa03C was plated at a density of 1 × 10□cells per 10-cm dish and allowed to adhere overnight. For each experimental condition, 3–10 dishes were used per biological sample to obtain sufficient material for downstream PTM proteomics analysis. Cells were treated with either DMSO (vehicle control for 120 min) or APX2014 (15 µM for 30, 60 or 120 min). The concentration selected for APX2014 is the 24 h IC_50_ from previously published work for these cells (13). The experiment was performed in four independent batches, each containing the complete time course (30, 60, and 120 min). At the indicated time points, cells were harvested by trypsinization, collected by centrifugation, and washed with cold PBS with 100 mM N-ethylmaleimide (NEM) to preserve redox-sensitive PTMs. Cell pellets were then snap-frozen and stored at −80°C until PTM proteomic analysis.

### Sample processing for multi-PTM proteomics

Cell pellets were lysed in ice-cold buffer containing 250 mM MES (pH 6), 5% SDS, 1% Triton X-100, 10 mM EDTA, 100 mM NEM, 10 mM nicotinamide, 10 mM sodium butyrate, 2 µM SAHA, 10 mM NaF, and 1% phosphatase inhibitor cocktails 2 and 3 (Sigma). Samples were incubated in the dark at room temperature for 30 min, briefly sonicated, clarified by centrifugation (13,000 rpm, 10 min, 4 °C), and the supernatants were incubated at 55 °C and 850 rpm for 30 min. Protein concentration was determined by BCA.

For SP3 cleanup and digestion, 500 µg protein per sample (adjusted to 1 µg/µL) was processed on a KingFisher Flex as described previously (18, 19). Ethanol was added to 50% (v/v) to bind proteins to mixed Sera-Mag SpeedBeads (E3 and E7, 5:1 of bead: protein), followed by three washes with 80% ethanol. Beads were transferred into 50 mM HEPES (pH 7.7) containing trypsin and LysC (1:100 each) and digested at 37°C, 850 rpm for 3 h; peptides were collected and combined with a bead rinse (250 mM HEPES, pH 7.7, 5% ACN). For labeling, 200 µg of peptides per sample were reacted with TMTpro 18-plex at a 2.5:1 (w/w) TMT:peptide ratio for 1 h at room temperature (850 rpm), then quenched with hydroxylamine. Labeled peptides were desalted using C18 SPE (Waters Sep-Pak, 50 mg) and split for global proteome quantification and PTM enrichments (cysteine oxidation, phosphorylation, acetylation).

### Enrichment of PTM peptides

Peptides containing oxidized cysteines were reduced with 5 mM DTT and enriched using a peptide-level resin-assisted capture (RAC) workflow as previously described (17, 20). The resulting cysteine-containing peptides were alkylated with IAM (iodoacetamide, at a final concentration 4:1 to DTT concentration in the elution), followed by C18 SPE cleanup and high-pH reverse phase capillary LC fractionation. In total, 12 fractions were collected for LC-MS/MS analysis of the redox proteome. For global protein abundance quantification, an aliquot of TMT-labeled peptides was alkylated with IAM, cleaned up by C18 SPE and fractionated by high-pH RPLC into 12 fractions in parallel with redox proteome samples.

For phosphopeptide enrichment, an aliquot of TMT-labeled peptides was first fractionated into 12 fractions by high-pH RPLC, then automated Immobilized Metal Affinity Chromatography (IMAC) was conducted on Agilent Bravo system as reported(21). The flowthroughs from IMAC enrichment (12 fractions) were collected and used for acetylation enrichment with the PTMScan HS Acetyl-Lysine Motif [Ac-K] kit (Cell Signaling Technologies #46784) on the KingFisher Flex, as described (18, 21).

### LC-MS/MS

All samples were analyzed on a Thermo Vanquish Neo UHPLC system coupled to an Orbitrap Exploris 480 mass spectrometer (Thermo Scientific). Peptides were separated on an in-house packed Acquity 1.7 μm BEH C18 (Waters) column (25 cm × 75 μm inner diameter) using 120 min gradient. The mobile phase A was 0.1% formic acid (FA) in water. Mobile phase B was acetonitrile containing 0.1% FA. Full MS scans were acquired in m/z range of 380 – 1800 with a resolution of 60K by data-dependent acquisition (DDA) mode. Standard automatic gain control (AGC) target was applied. Maximum injection time was set to Auto. The top 20 most intense precursor ions from full MS scans were selected (with 0.7 m/z isolation window) and fragmented using higher-energy collision dissociation (HCD). Dynamic exclusion duration was 45 s. MS/MS scans were acquired at a resolution of 45K with AGC target of 250%. The first mass was set as 120 m/z to detect TMT reporter ions.

### Data analysis

Raw MS data were analyzed by MS-GF+ (22) against the *Homo sapiens* UniProt protein database (released 2023-09). A partially tryptic search was conducted with precursor tolerance of 20 ppm. Static modifications were TMT on lysine and peptide N-terminal (+304.2071 Da). For global and redox proteomics samples, dynamic modifications included oxidation on methionine (+15.9949 Da), alkylation of cysteine with NEM (+125.0477 Da), and cysteine carbamidomethylation (+57.0215 Da). For phosphoproteomics samples, dynamic modifications were phosphorylation of serine, threonine, and tyrosine (+79.9663 Da), methionine oxidation, and cysteine alkylation with NEM. For acetylatomics samples, besides methionine oxidation and cysteine alkylation, dynamic modifications also included a −262.1966 Da on lysine, corresponding to the mass difference between a TMT tag and an acetyl group.

The peptide spectrum matches (PSMs) from MS-GF+ were filtered to absolute delta precursor mass < 10 ppm and peptide Q-value < 0.01 to obtain a false discovery rate (FDR) at the peptide level < 1%. Contaminant and decoy PSMs were removed. Reporter ion intensities of TMT18 were extracted using MASIC(23). These initial data processing steps were performed using R package PlexedPiper (https://github.com/PNNL-Comp-Mass-Spec/PlexedPiper).

For global proteomics, the TMT reporter intensity protein matrix was first filtered to proteins quantified across all samples. Intensities were then normalized using the global median method and subsequently batch-corrected using an empirical Bayes approach (ComBat), with PCA used to confirm removal of batch effect before downstream statistical analyses (24). For PTM datasets, peptide-level PTM crosstabs were first collapsed to modification site level by grouping entries that mapped to the same protein residue. The roll-up also retained site multiplicity information (i.e., the number of modified peptides contributing to a site). Site-level PTM intensities were then normalized in a global-proteome–referenced manner (i.e., adjusted relative to the corresponding global protein abundance) and batch-corrected with ComBat. Differential abundance was assessed by pairwise comparisons to the control condition using two-sample t-tests with Benjamini–Hochberg multiple-testing correction.

### Bioinformatics

Significant proteins and PTM sites were defined using both adjusted p-value and fold-change cutoffs. For functional interpretation, significant protein/site lists were tested for Gene Ontology and Reactome pathway enrichment using clusterProfiler. For structural analyses, AlphaFold2-predicted protein structures (https://alphafold.ebi.ac.uk/; model_v6) were retrieved using UniProt accession and parsed in R using bio3d (25). PTM sites of interest were mapped to the corresponding residues on the specified protein chain. PTM locations were visualized in 3D using r3dmol, rendering a cartoon representation of each protein and marking PTM sites as colored spheres (yellow for oxidized cysteines, red for Ser/Thr/Tyr phosphorylation, and blue for lysine acetylation). To assess spatial relationships among PTM sites, pairwise Cα–Cα Euclidean distances (Å) were computed and visualized using ComplexHeatmap.

### Mitochondrial assay (substrate consumption)

Mitochondrial substrate utilization was assessed using S-1 Mitoplates (Biolog, Hayward, CA, cat#14105) according to the manufacturer’s instructions. Prior to the assay, wells were preactivated with the supplied Assay Mix (Biolog 6X Redox Dye MC, cat#74353) for 1 h at 37°C to ensure substrate solubilization. Pa03C cells were treated with either DMSO vehicle or 15 µM APX2014 for 24 h. Following treatment, cells were detached with trypsin, pelleted by centrifugation, and counted using trypan blue exclusion. Cells were then resuspended in the assay buffer (Biolog 2X Mitochondrial Assay Solution, cat#72303) provided by the manufacturer and seeded into Mitoplates at a density of 5 × 10^4^ cells per well. Immediately after plating, absorbance at 590 nm was recorded at 5 min intervals over a 4 h incubation period at 37°C. Data analysis was carried out using GraphPad Prism 11. Differences in average reaction rates were evaluated using two-way ANOVA, while changes in kinetic slopes were analyzed by linear regression. Statistical significance was defined as p < 0.05.

### Immunofluorescence and measurement of mitochondria volume

Pa03C cells were seeded (3 × 10^5^) per well on cover slips in 6-well plates. After treatment with Ref-1 inhibitor for 30, 60, 120 min, and 24 h with 15μM APX2014, cells were fixed with paraformaldehyde-Triton X-100 fixation buffer (4% methanol-free paraformaldehyde, and 0.5% Triton X-100 in PBS) at room temperature for 10 minutes. Fixed cells were washed twice with wash buffer (0.03% Triton in PBS) X-100 then permeabilized with blocking buffer (5% BSA, 0.5% Triton X-100, and 0.05% SDS in PBS) for 30-60 minutes. Cells were incubated in primary anti-mitochondria (Millipore Sigma MAB1273) and anti-Ref-1 (Novus NB100-101) antibodies diluted 1:100 in wash buffer, either at room temperature for at least one hour or overnight at 4°C. Coverslips were then washed with wash buffer three times and incubated in fluorophore-conjugated secondary antibodies (Invitrogen; diluted 1:2,000 in wash buffer) at room temperature for one hour. Coverslips were then washed with wash buffer three times, rinsed once in distilled water, then mounted onto slides in Prolong Diamond Antifade Mountant with 4’,6-diamidino-2-phenylindole (Invitrogen). Slides were cured at least 24 hours at room temperature before imaging. Images were acquired with a Deltavision Ultra microscope using a 100x objective. Images were acquired in z-series with sections of 0.2µm each and deconvolved using SoftWorx software (Leica). Mitochondria volume was measured using the surfaces function in Imaris software (Bitplane). All images were processed using Imaris.

## Statistical Analysis

Differential protein abundance and PTM site analyses were performed by pairwise comparison of APX2014-treated and DMSO-treated samples using two-sample t-tests with Benjamini–Hochberg false discovery rate (FDR) correction for multiple testing. Principal component analysis, hierarchical clustering, UpSet analyses, pathway enrichment, and structural mapping were performed in R. Functional enrichment analyses were conducted using clusterProfiler. Mitochondrial substrate utilization assays were analyzed using GraphPad Prism 11. Average reaction rates were compared using two-way ANOVA, and kinetic differences were evaluated by linear regression. Mitochondrial immunofluorescence data were analyzed using Welch’s t test. Unless otherwise indicated, P < 0.05 or FDR-adjusted P < 0.05 was considered statistically significant.

## Results

### Short-term APX2014 exposure causes significant PTM changes in Pa03C cells

To investigate the systems-level impact and underlying molecular mechanism of the Ref-1 inhibitor APX2014, pancreatic cancer cell line, Pa03C were treated with APX2014 for 30, 60, and 120 min alongside DMSO vehicle control. Samples following treatment were multiplexed using TMTpro 18-plex and analyzed using an integrative multi-PTM proteomics workflow to quantify global protein abundance and PTMs, including cysteine oxidation, phosphorylation, and acetylation **(Fig. 1A; Supplementary Data 1)**. Across all conditions, we identified 22,255 oxidized (redox) peptides corresponding to 18,183 cysteine sites **(Fig. 1B)**; most carried a single oxidation event (14,122 sites), with fewer containing two (3,240) or three or more sites (821) (**Fig. S1A**). We also quantified 45,359 phosphopeptides corresponding to 40,831 S/T/Y sites, which were dominated by singly phosphorylated sites (30,451), with 8,930 doubly and 1,450 triply (or more) sites **(Fig. S1B)**. Phosphorylation occurred predominantly on serine (32,536 sites) and threonine (6,875 sites), with fewer on tyrosine (1,420 sites) (**Fig. S1C**). For acetylation, we identified 23,441 acetylated peptides mapping to 14,120 acetylation sites, with the vast majority being singly acetylated (13,569 peptides), and fewer containing two (436) or three or more (115) acetylation sites (**Fig. 1B and Fig. S1D**).

**Figure 1.**
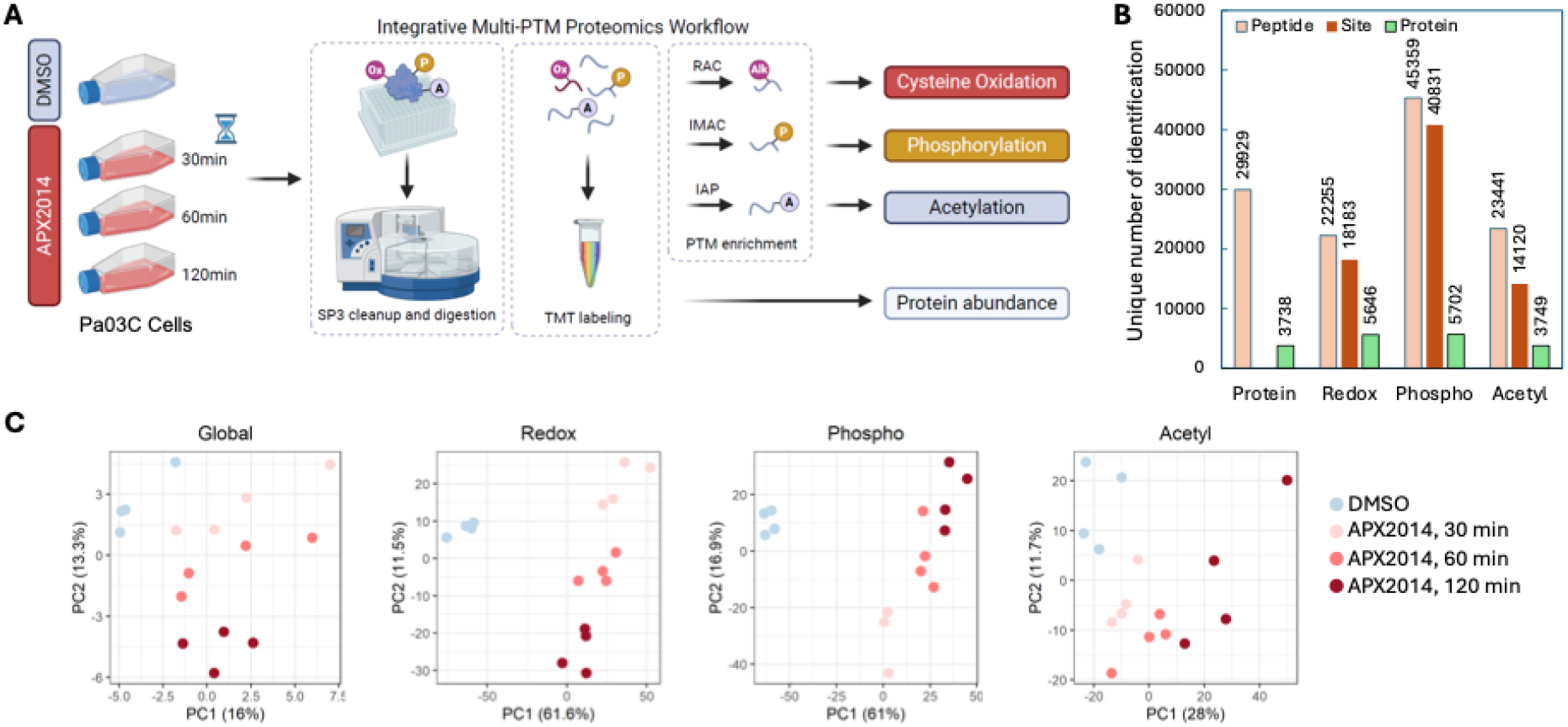
Integrative multi□PTM proteomics profiling of acute APX2014 response in pancreatic cancer cells. (**A**) Pa03C cells were treated with DMSO or APX2014 for 30, 60, or 120 min and processed using an SP3-based workflow with TMT labeling, followed by parallel enrichment and LC–MS/MS analysis to quantify global protein abundance and three PTM layers: cysteine oxidation (via resin-assisted capture, or RAC), phosphorylation (immobilized metal affinity chromatography, or IMAC), and acetylation (immunoaffinity purification, or IAP). (**B**) Bar plot showing modified peptides, sites, and proteins. (**C**) PCA showing the separation patterns among experimental groups.

Principal component analysis (PCA) conducted separately for each dataset showed clear separation between DMSO-and APX2014-treated samples, with the most significant separation seen in the redox and phosphorylation datasets, with clear time dependence (**Fig. 1C**). Overall, these results demonstrate that APX2014 triggers rapid, system-wide remodeling, primarily evident as changes in cysteine oxidation and phosphorylation, which align with early redox and signaling disturbances before broader shifts in protein abundance.

### PTM regulation on proteins associated with APX2014

To assess the importance of the PTM modifications in the APX2014-induced cellular response, we examined the PTM levels of proteins previously shown to be involved in Ref-1–impacted signaling nodes (26, 27). The list consisted of 20 proteins, and all 20 showed statistical significance in at least one PTM in our dataset (**Table 1**). Assessment of site-resolved PTM dynamics of these proteins across the acute APX2014 time course revealed many responsive sites to Ref-1 inhibition. For example, we observed broad coverage on 12 oxidation sites, 7 phosphorylation sites, and 6 lysine acetylation sites for NF□κB1 (**Fig. 2A**). Clustering of these PTMs showed that many oxidation sites (C61/C261/C272) and acetylation sites (K76/K78/K79/K243) share similar temporal pattern of change to APX2014 treatment and they are structurally located in the N□ terminal Rel homology region (RHD) (**Fig. 2B**). In addition, another cluster of PTMs were found in its C□terminal (C925; S893/S907/S923/S927/S937/T939/S947; and Lys K830). Many sites are spatially in close proximity to the well-known redox-sensitive site C61 in the RHD (**Fig. S2A**) (28). Thus, these data indicate that Ref-1 drives coordinated PTM remodeling in the DNA-binding region to control NF□κB1 activity.

**Figure 2.**
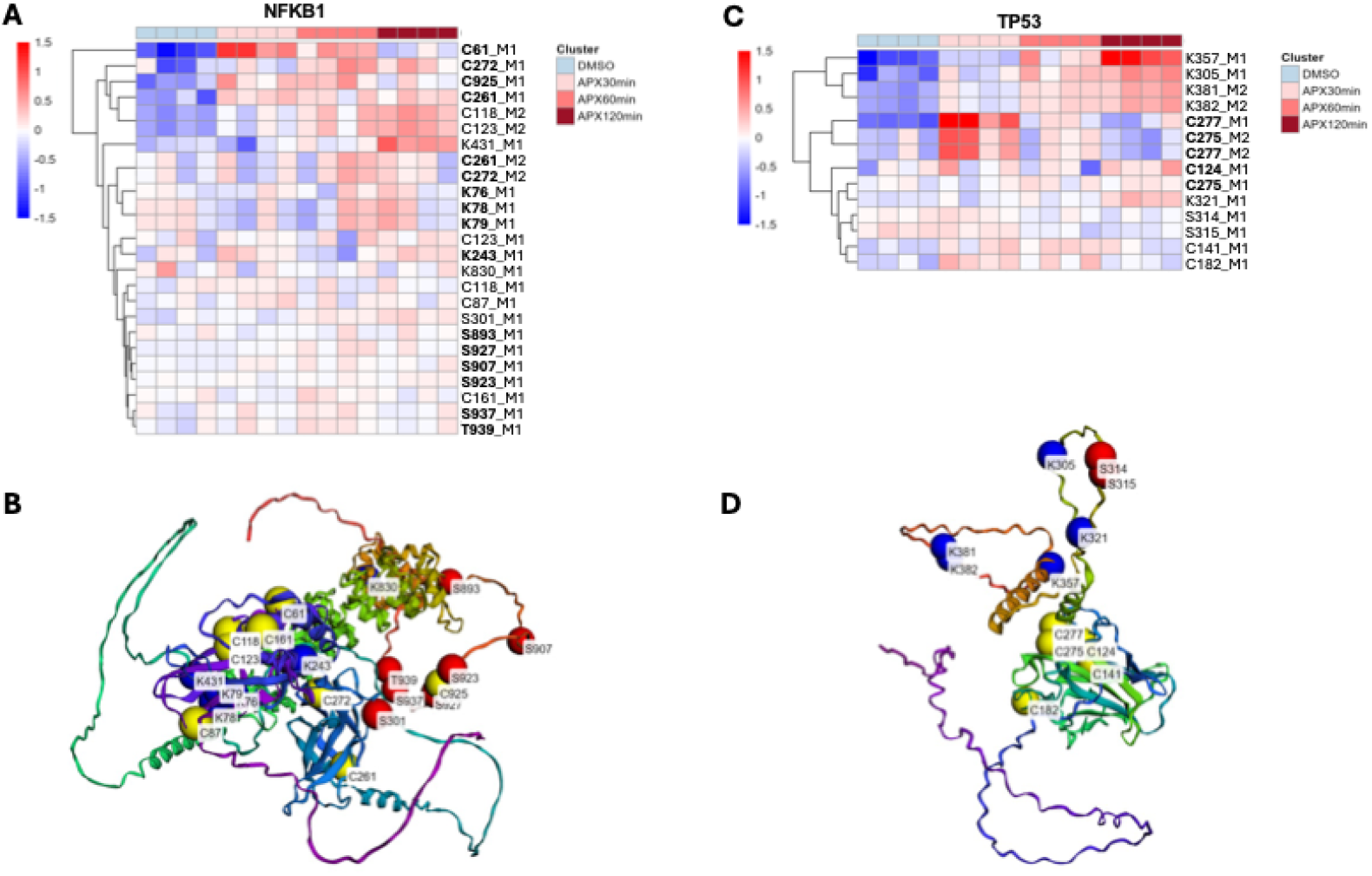
**APX2014 induces coordinated PTM remodeling on NF-**κ**B1 and p53.** (**A**) Heatmap of site-resolved PTMs quantified on NF-κB1 across the acute APX2014 time course (DMSO, 30, 60, and 120 min). (**B**) Structural mapping of quantified NF-κB1 PTM sites onto a protein structure model. (**C**) Heatmap of PTMs for p53 (TP53). (**D**) Structural mapping of p53 PTM sites. In heatmaps, values are shown as normalized site-level intensities (row-scaled), and sites are clustered hierarchically to highlight shared temporal trajectories. Bolded residues are those mentioned in the text. In structural models, the protein backbone is shown as a cartoon representation, and PTM sites are highlighted as spheres (yellow, cysteine oxidation; red, phosphorylation; blue, lysine acetylation).

**Table 1:**
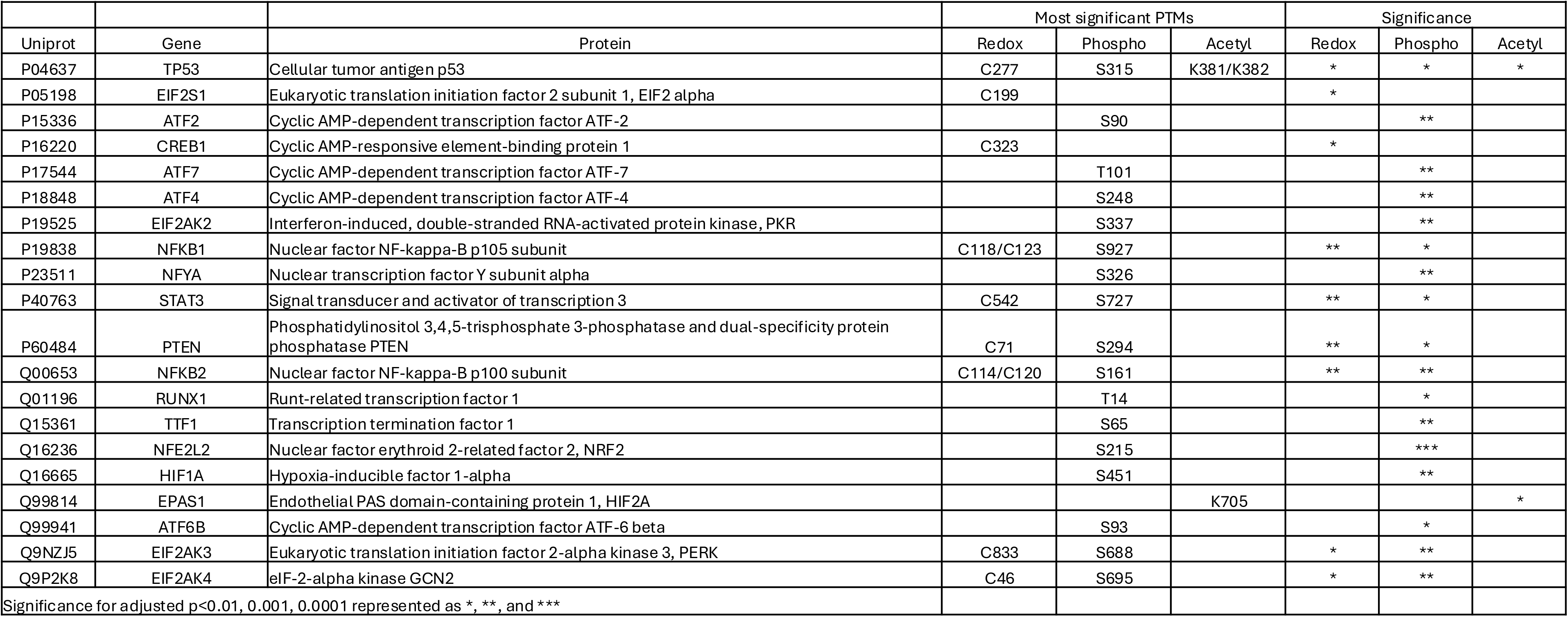
List of significant previously known Ref-1 regulated PTM modifications.

For p53, we observed significant increase in oxidation of C275/C277 at 30 min, and increased acetylation at K357/K305/K381/K382 at 60 and 120 min (**Fig. 2C**). Structural mapping onto the AlphaFold model localized multiple quantified cysteines within the p53 DNA-binding core (including C124 and C275/C277), whereas the regulated lysine acetylation and serine phosphorylation sites mapped primarily to more flexible regulatory regions outside the core (**Fig. 2D**). Consistent with this organization, distance mapping indicated that several cysteine sites form a spatially proximal cluster (e.g., within the DNA-binding domain; **Fig. S2B**).

### APX2014-induced PTM signatures in Pa03C cells

We next performed differential analysis at each timepoint by comparing APX2014-treated cells to the matched DMSO controls within each omics layer. Consistent with the PCA analysis, APX2014 induced minimal changes in global protein abundance during the 30–120 min time window (**Fig. S3**), whereas PTM layers showed rapid, widespread remodeling (**Fig. 3A-C**). In particular, cysteine oxidation showed a strong, predominantly increased response, with 2,490 significantly more oxidized Cys sites and only 1 more reduced sites, respectively, at 30 min (**Fig. 3A; Supplementary Data 2)**. The shift toward to a more oxidized proteome persisted to 60 and 120 min, although the number of significantly more oxidized Cys sites decreased compared to 30 min. Phosphorylation changes were also extensive and time-dependent, with both significantly increased and decreased sites observed (**Fig. 3B; Supplementary Data 3)**. In contrast to redox regulation, the number of significant phosphorylation sites showed an increase with time, with 532, 1,146, and 1,683 increasing sites at 30, 60, and 120 min, respectively. Acetylation changes were comparatively modest at early time points but became more pronounced by 120 min, indicating a delayed response relative to redox and phosphorylation (**Fig. 3C; Supplementary Data 4)**.

**Figure 3.**
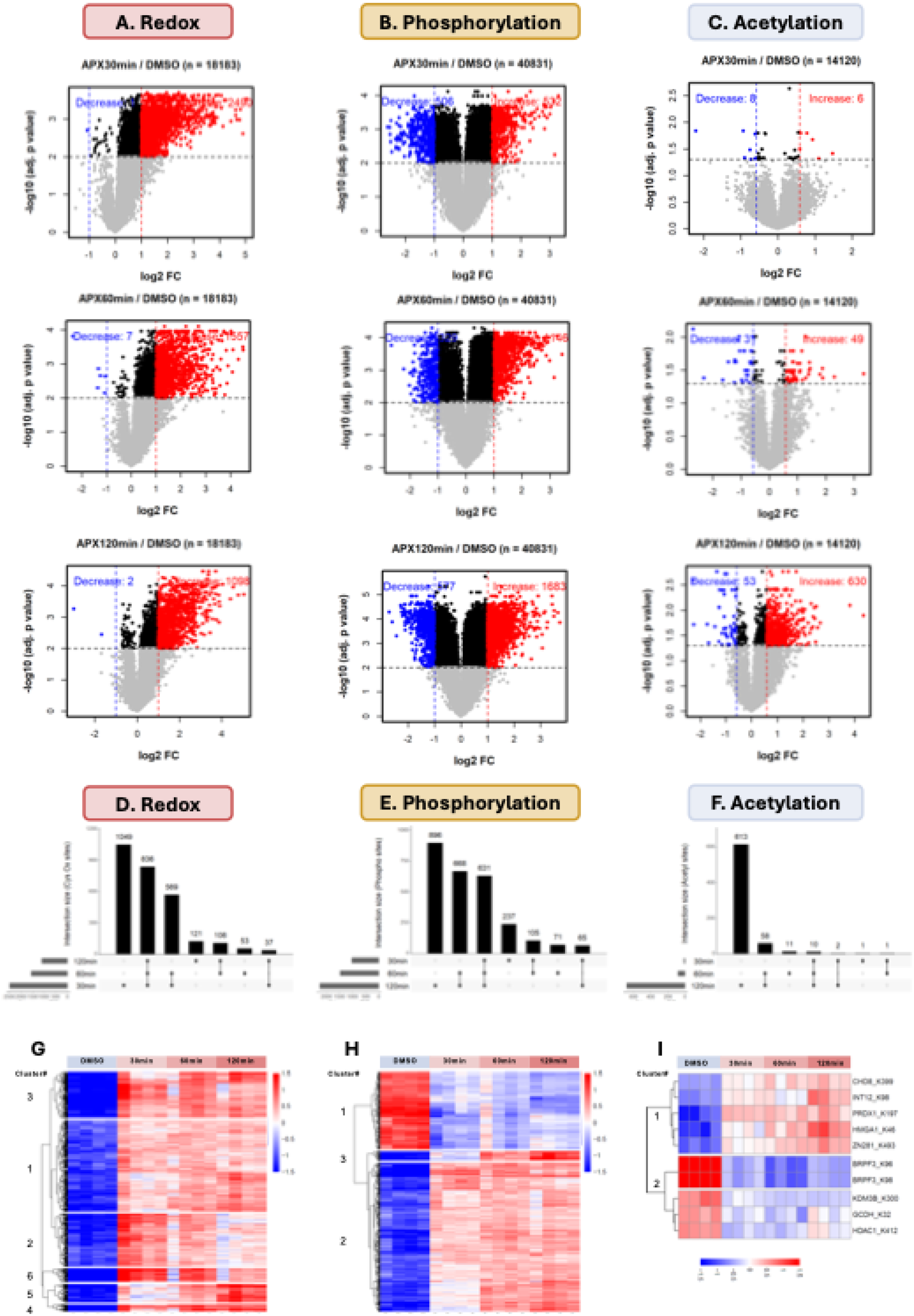
PTM temporal dynamics by APX2014 treatment. Volcano plots for (**A**) cysteine oxidation (redox) sites; (**B**) phosphorylation (S/T/Y) sites and; (**C**) lysine acetylation sites. Points are colored to indicate significantly decreased (blue) or increased (red) features, with non-significant features shown in gray/black; dashed lines denote fold-change and adjusted p-value thresholds. The number of quantified features (n) is indicated for each comparison. (**D–F**) UpSet plots showing the overlap of significantly changed sites (APX2014 vs DMSO) across 30, 60, and 120 min for cysteine oxidation (redox, **D**), phosphorylation (**E**), and acetylation (**F**), respectively. (**G–I**) Heatmaps showing hierarchical clustering of sites that were significantly regulated at all three time points within each PTM type. For each PTM site (row), levels were median-centered across all samples, so that red and blue indicate levels above and below the median, respectively.

To evaluate the temporal consistency of APX2014-regulated PTM changes, we summarized significant sites across 30, 60, and 120 min using UpSet analysis (**Fig. 3D–F**). Across the three time points, the redox dataset contained 836 shared cysteine sites (**Fig. 3D)**, and the phosphorylation dataset contained 631 shared phosphorylation sites (**Fig. 3E),** indicating sustained, significant PTM remodeling throughout the acute response window. By contrast, significant acetylation sites were mostly detected at 120 min with only 10 common sites across time points (**Fig. 3F)**. We then focused on the subset of overlapping significant sites detected at all three time points and visualized their dynamics by hierarchical clustering (**Fig. 3G–I**). These shared site heatmaps highlight several temporal trajectories for cysteine oxidation and phosphorylation across the full treatment course, including a cluster of cysteine sites that show clear shifts already apparent at 30 min (Cluster 5, **Fig. 3G**). Cluster 5 consisted of 191 cysteine sites on 170 proteins (**Supplementary Data 5**), and enrichment of this early-responding set is strongly biased toward mitochondrial (mitochondrial organization and gene expression/translation as well as mitotic nuclear division and chromosome segregation) and cell-cycle pathways (**Fig. S4A**). Consistent with these processes, cellular component enrichment highlighted mitochondrial structures (mitochondrion, mitochondrial nucleoid, organellar ribosome) alongside nuclear/mitotic features (nuclear envelope/membrane, spindle, and condensed chromosomes; **Fig. S4B**).

In contrast to one-sided shift in cysteine oxidation, both up-and down-regulation were found in the shared sites for phosphorylation (**Fig. 3H**) and acetylation (**Fig. 3I**). Reactome analysis further indicated that the bidirectional phosphoproteomic response involved distinct biological processes. Proteins containing increased phosphorylation sites (423 phosphorylation sites mapping to 310 proteins) were enriched in RHO GTPase signaling, pre-mRNA processing, SUMOylation, and cellular responses to heat stress (**Fig. S5A**). In contrast, proteins containing decreased phosphorylation sites (208 sites mapping to 154 proteins) were predominantly associated with mitotic progression, including M phase, chromosome segregation, mitotic spindle formation, and epigenetic regulation of gene expression (**Fig. S5B**). In the acetylome, APX2014 similarly elicited a bidirectional response, with increased acetylation at sites including CHD8-K399, INT12-K98, PRDX1-K197, HMGA1-K46, and ZN281-K493, particularly at later time points. Conversely, acetylation of BRPF3-K96/K98, KDM3B-K300, GCDH-K32, and HDAC1-K412, decreased following APX2014 treatment (**Fig. 3I**).

### Pathway enrichment reveals early mitochondrial and translational remodeling by APX2014

To contextualize APX2014-driven changes at the pathway level, we performed enrichment analysis using significantly regulated PTM sites/proteins. For the redox, phosphorylation, and acetylation datasets, the top Reactome pathways (**Fig. 4A,C,E**) and top GO cellular component terms (**Fig. 4B,D,F**) are shown. The results showed that the three PTM layers exhibit distinct regulation following Ref-1 inhibition with APX2014. First, the PTMs related to redox revealed a pronounced mitochondrial signature, with the enrichment of pathways linked to mitochondrial translation (initiation/elongation/termination), mitochondrial protein degradation, respiratory electron transport, and central carbon metabolism (e.g., TCA cycle, **Fig. 4A**). Consistently, GO cellular component enrichment highlighted ribosome/large ribosomal subunit, mitochondrial ribosome, and mitochondrial protein-containing complexes (**Fig. 4B**). Together, these results indicate that APX2014 rapidly perturbs oxidation states of proteins associated with mitochondrial gene expression and oxidative phosphorylation, pointing to mitochondria as a prominent early redox-responsive compartment.

**Figure 4.**
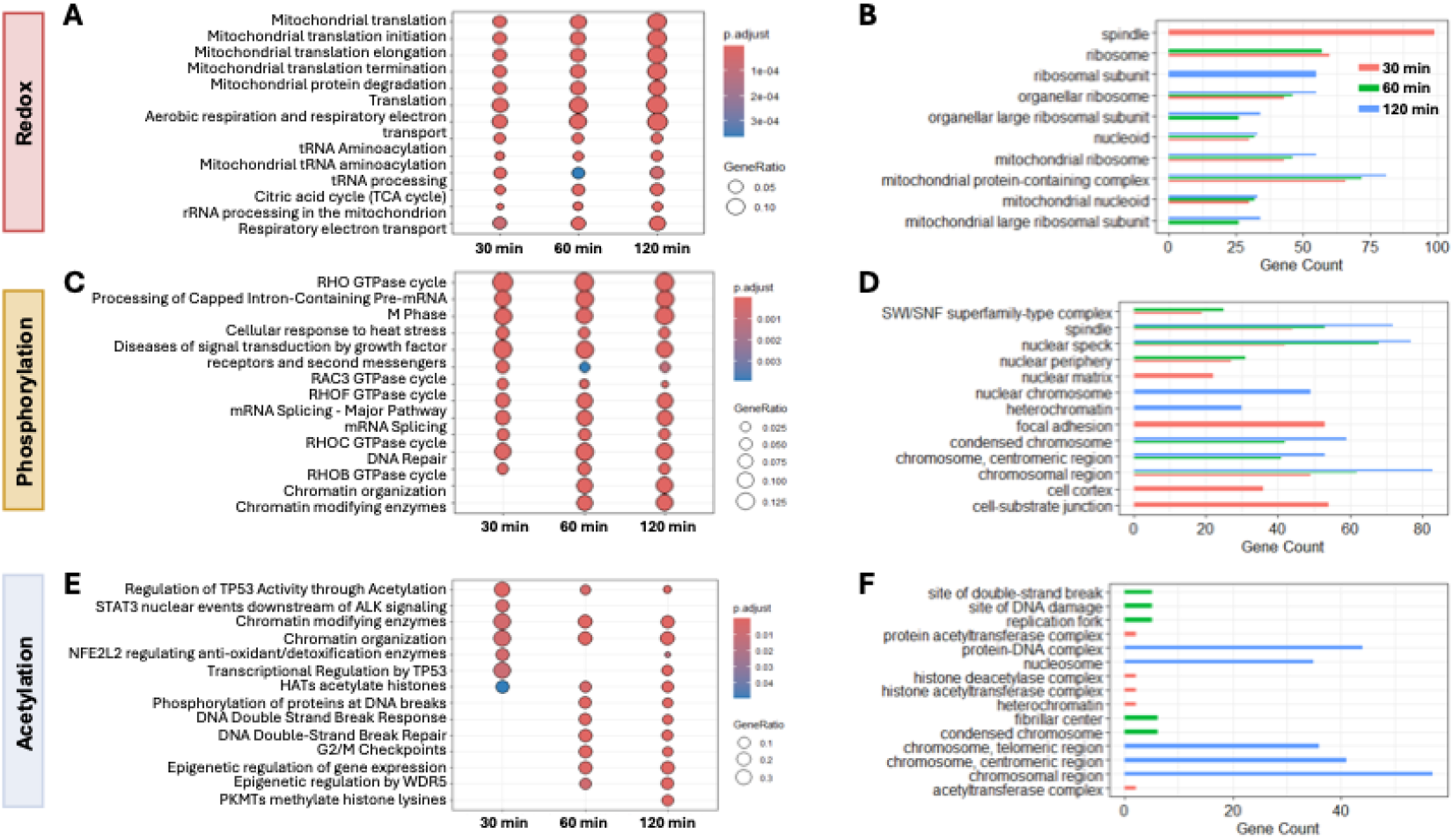
Pathway and subcellular enrichment of APX2014-regulated PTM sites. Reactome pathway enrichment of proteins containing significantly regulated sites is shown for (**A**) cysteine oxidation (redox), **(C)** phosphorylation, and (**E**) acetylation across 30, 60, and 120 min APX2014 treatment; dot size reflects gene ratio and color indicates Benjamini–Hochberg adjusted p value. Corresponding GO Cellular Component enrichments are shown for (**B**) redox, (**D**) phosphorylation, and (**F**) acetylation, displayed as gene counts for the top-enriched terms at each time point.

In contrast, phosphorylation events preferentially captured signaling and structural regulation centered on nuclear architecture and cell cycle machinery. Reactome pathways were enriched for Rho-family GTPase cycles, mRNA splicing, DNA repair, and chromatin organization/modifying enzymes (**Fig. 4C**), consistent with kinase-driven remodeling of cytoskeletal/signaling pathways, nuclear RNA processing and genome maintenance responses. The phosphoregulated proteins were localized to spindle, nuclear speck, chromosome/heterochromatin, and focal adhesion/cell–substrate junction (**Fig. 4D**), suggesting that APX2014 rapidly engages phosphosignaling networks governing mitotic regulation, chromatin state, and cell adhesion-related signaling.

Finally, the acetylation sites were enriched in nuclear/chromatin terms such as TP53 regulation (via acetylation), HAT/chromatin-modifying enzymes, and epigenetic regulation, alongside DNA double-strand break response/repair (**Fig. 4E**). Consistently, GO Cellular Component terms highlighted nucleosome/histone and protein acetyltransferase complexes and chromosomal regions (**Fig. 4F**), indicating that Ref-1-responsive acetylation primarily reflects chromatin-and transcription-associated stress/DNA damage pathways. Thus, each PTM layer provided distinct insights: redox primarily mapped to mitochondrial translation/respiration, phosphorylation to Rho GTPase signaling, splicing, and mitotic/adhesion structures, and acetylation to histone/chromatin machinery and p53-linked transcriptional control. Together, the multi-PTM data suggest an early mitochondrial redox response by APX2014, accompanied by rapid phosphosignaling rewiring and broader chromatin/transcription changes reflected in acetylation.

### Cell-based validation confirms impact of Ref-1 inhibition on mitochondrial function predicted by PTM proteomics

To validate the mitochondrial alterations suggested by PTM proteomic analyses, we performed two assays to assess the changes in the mitochondria: a plate-based mitochondrial substrate utilization assay and quantitation of mitochondrial volume following treatment with APX2014. The plate-based assay measures the ability of intact cells to metabolize individual mitochondrial substrates and provides a functional assessment of mitochondrial activity (13, 14). APX2014 treatment significantly impaired utilization of the TCA cycle substrates α-ketoglutarate, succinate, fumarate, and malate, resulting in a 58– 61% reduction in average reaction rate compared with DMSO-treated controls (**Fig. 5A,B**). Consistent with these findings, kinetic analyses demonstrated reduced substrate utilization throughout the assay period, with endpoint reductions ranging from approximately 40–51% (**Fig. 5C**). Although additional metabolic alterations were observed across the broader substrate panel (**Fig. S6**), the most pronounced effects were detected in TCA cycle-associated substrates, which were selected for further analysis.

**Figure 5.**
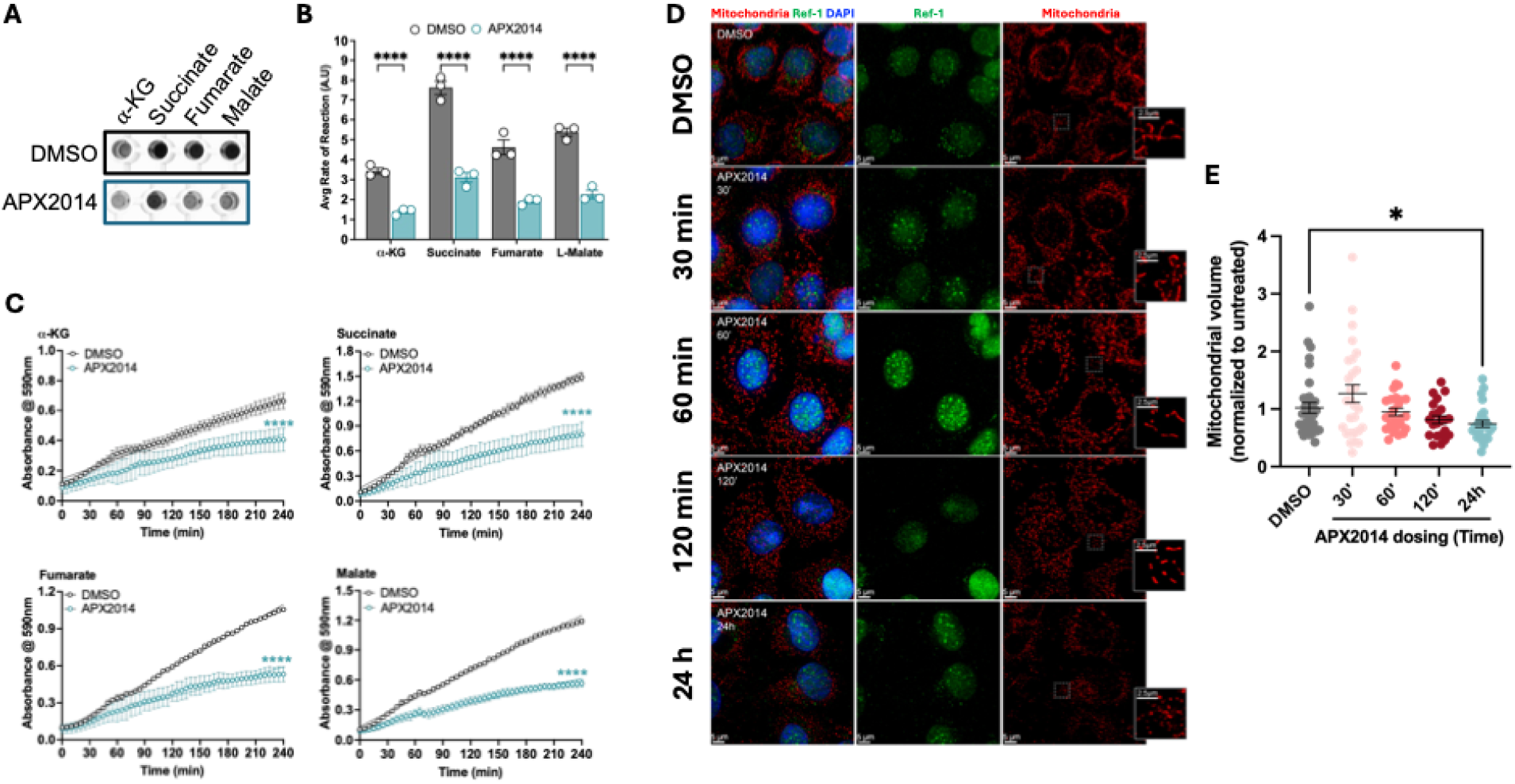
Ref-1 inhibition by APX2014 decreases mitochondrial TCA-cycle substrate utilization and mitochondrial volume in Pa03C cells. (**A**) Representative colorimetric assay images showing the utilization of tricarboxylic acid (TCA) cycle metabolites in Pa03C cells treated with vehicle (DMSO) or APX2014 (15 μM) for 24 h. Wells correspond to α-ketoglutarate (α-KG), succinate, fumarate, and malate substrates. (**B**) Quantification of average reaction rates for individual TCA cycle metabolites (arbitrary units, A.U). Rates were calculated from the initial 90 min of the 240 min assay. Data are presented as mean ± SE from n = 3 independent biological replicates. Statistical significance was determined using two-way ANOVA. ****p< 0.0001. (**C**) Kinetic analysis of substrate utilization over time demonstrating decreased metabolic activity in APX2014-treated cells relative to vehicle-treated cells for α-KG, succinate, fumarate, and malate. Data are presented as mean ± SE from n = 3 independent biological replicates. Statistical significance between the slopes was determined using simple linear regression. ****p< 0.0001. **(D)** Representative immunofluorescence images of mitochondria (red) and Ref-1 (green) co-staining. DAPI was used for nuclear staining. (**E**) Quantification of average mitochondrial volume per cell following APX2014 treatment relative to DMSO control. Data are pooled from two independent experiments. P value was calculated by Welch’s t test (p=0.0136).

To further confirm the PTM data that Ref-1 inhibition alters mitochondrial function and potentially morphology, mitochondrial volume was quantified by immunofluorescence (IF) following treatment with APX2014 across early and late time points. Ref-1 inhibition did not significantly alter mitochondrial volume at early time points, including 30, 60, and 120 minutes after treatment (**Fig. 5D,E**), suggesting that the PTMs caused by Ref-1 inhibition do not immediately affect mitochondrial morphology. In contrast, the 24 h time point showed significantly reduced mitochondrial volume (p=0.0136), indicating that prolonged Ref-1 inhibition leads to a measurable decrease in mitochondrial volume and function (**Fig. 5D,E**). Taking these findings together suggests that the effects of Ref-1 inhibition on mitochondrial function are time-dependent and emerge after the initial early events that were captured through the PTMs. These functional data support our PTM proteomic findings and are consistent with our and others previous transcriptomic studies demonstrating that Ref-1 redox signaling regulates pathways critical for mitochondrial function (14, 29).

### Rapid redox regulation by APX2014

To define downstream consequences of Ref-1 redox signaling, we further analyzed proteins that showed rapid oxidation after APX2014 treatment, represented by cluster 5 in **Fig. 3**. Term–gene network analysis organized these proteins into interconnected modules dominated by mitochondrial function and mitotic regulation, with additional smaller clusters linked to cellular redox homeostasis, RNA modification, and fatty-acid β oxidation (**Fig. 6**). This cluster includes well-established redox-responsive proteins such as PRDX1 and PRDX4 (peroxiredoxins) and TXN2 (mitochondrial thioredoxin), all of which contain catalytic cysteines that undergo rapid, reversible oxidation during peroxide detoxification and redox signaling (30–32). Consistent with these findings, our group recently demonstrated that PRDX1 expression modulates cellular response to Ref-1 inhibition, as PRDX1-deficient pancreatic cancer cells are exquisitely sensitive to APX2014 both *in vitro* and *in vivo* (11). Cluster 5 also included NNT (nicotinamide nucleotide transhydrogenase), a key mitochondrial enzyme that sustains NADPH pools required for thioredoxin-and glutathione-dependent reduction capacity and is often linked to mitochondrial redox buffering. Importantly, prior work with an additional second-generation Ref-1 inhibitor, APX2009 demonstrated downregulation of NNT in pancreatic cancer cells following treatment (14). Together, these data indicate that Ref-1 redox inhibition rapidly disrupts mitochondrial redox control and suggest that early cysteine oxidation events may precede broader transcriptional regulation of mitochondrial stress-response pathways.

**Figure 6.**
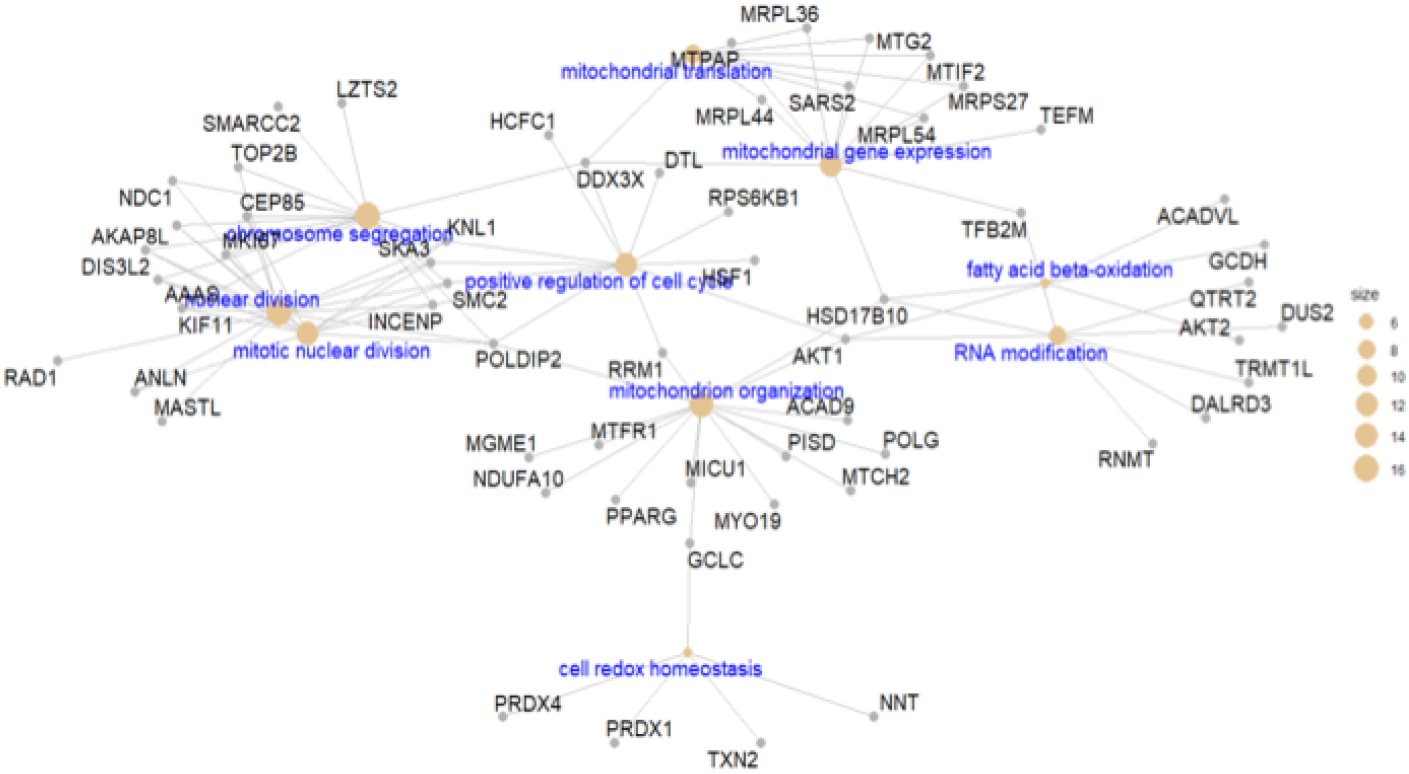
Proteins exhibiting rapid redox-dependent changes after APX2014 treatment. Term–gene network linking significantly enriched GO terms (blue labels) to the corresponding proteins (gray nodes); node size reflects the number of connections/annotated proteins per term.

## Discussion

Ref-1 is an important regulator of redox-sensitive transcriptional signaling and has been implicated in pancreatic cancer progression, stress adaptation, metabolism, and treatment resistance (10, 11, 13, 14, 26). Although previous studies have shown that pharmacologic Ref-1 inhibition can suppress tumor growth and alter metabolic and transcriptional pathways, the early biochemical events that precede these downstream effects remain incompletely defined (33, 34). To address this, we used the selective Ref-1 redox inhibitor APX2014 as a pharmacological tool and applied an integrated multi-PTM proteomics approach to quantify cysteine oxidation, phosphorylation, and lysine acetylation together with global protein abundance over an acute 30–120 min treatment period in PDAC cells. Ref-1 inhibition produced substantial PTM remodeling while overall protein abundance remained largely stable, indicating that the initial response is driven primarily by changes in protein regulatory states rather than protein quantity. Redox and phosphorylation profiles showed the clearest early separation between treated and control samples, consistent with rapid effects on cysteine oxidation and kinase-regulated signaling, whereas acetylation changes were more evident at later time points. Together, these findings suggest a temporally organized response in which early redox and phosphorylation changes are followed by broader acetylation-associated remodeling involving mitochondrial, chromatin-associated, stress-response, and cell-cycle pathways.

The response to Ref-1 inhibition followed a clear temporal hierarchy. Cysteine oxidation was the earliest and most sustained response, phosphorylation changes were also widespread and time dependent, and acetylation became more prominent at later time points (**Fig. 3**). This pattern fits with the known role of Ref-1 in maintaining redox-sensitive proteins in a reduced and functional state (35–37). Blocking Ref-1 redox activity would therefore be expected to alter reactive cysteine residues rapidly, with secondary consequences for redox-sensitive kinases, phosphatases, protein interactions, and chromatin-associated regulatory processes. The large number of redox and phosphorylation sites that remained significantly changed across 30, 60, and 120 min, together with the stronger acetylation response at 120 min, supports a model in which Ref-1 inhibition first disrupts redox homeostasis and then affects additional PTM networks. These observations do not establish that oxidation directly causes every subsequent phosphorylation or acetylation event, but they define a temporal framework in which redox disruption precedes and may shape broader signaling adaptation.

A prominent early target that emerged out of this response is mitochondrial biology. Proteins with regulated cysteine oxidation were strongly enriched for mitochondrial translation, mitochondrial ribosomal function, respiratory electron transport, TCA-cycle metabolism, mitochondrial protein degradation, and redox homeostasis. The cellular-component analysis similarly identified mitochondrial ribosomes, nucleoids, and mitochondrial protein-containing complexes. These findings indicate that mitochondria are not merely affected after prolonged loss of cell viability but represent an early redox-responsive compartment following Ref-1 inhibition. This is particularly relevant in PDAC, where mitochondrial flexibility and redox buffering support survival under hypoxia, nutrient limitation, and therapeutic stress (38, 39).

The rapid regulation of PRDX1, PRDX4, TXN2, and NNT within the early redox-responsive network further implicates disruption of mitochondrial and cellular antioxidant systems. PRDX1 and PRDX4 use catalytic cysteines to reduce peroxides and can function both as peroxide scavengers and redox sensors (40). TXN2 provides reducing capacity within mitochondria, whereas NNT supports mitochondrial NADPH production required for thioredoxin-and glutathione-dependent antioxidant systems (41–43). Their coordinated oxidation is consistent with perturbation of the NADPH–thioredoxin– peroxiredoxin axis following Ref-1 inhibition. This interpretation also aligns with our recent findings showing that loss of PRDX1 sensitizes pancreatic cancer cells to Ref-1 inhibition (11). However, because the present dataset measures site-level oxidation rather than direct enzymatic activity or peroxide flux, these changes should be interpreted as evidence of altered redox regulation rather than definitive proof that each antioxidant pathway is functionally inactivated. Further, the functional mitochondrial data support the biological significance of the proteomic findings. APX2014 markedly reduced utilization of α-ketoglutarate, succinate, fumarate, and malate, with lower reaction rates and altered kinetics across all four TCA-cycle substrates. These findings demonstrate that the mitochondrial PTM signature is accompanied by measurable impairment of oxidative substrate utilization. The Mitoplate assay was performed after 24 h of treatment, whereas proteomics captured the first 2 h. Hence, the two datasets were not interpreted as simultaneous measurements, but rather their consistency supports a temporal model in which early PTM remodeling of mitochondrial and redox-regulatory proteins precedes later loss of mitochondrial metabolic capacity. This was supported by the mito-volume measurement data where changes in oxidation and phosphorylation at 30, 60, and 120 min are not phenotypically quantifiable on the mito-volume until the later timepoint of 24 h. Together with previous observations that Ref-1 inhibition decreases oxidative phosphorylation and TCA cycle substrate utilization, the current data further position mitochondria as an important downstream effector of Ref-1 redox signaling (14).

The phospho-proteomic analysis provided a complementary view of the early response to Ref-1 inhibition, revealing rapid remodeling of signaling networks involved in Rho-family GTPase signaling, mRNA processing, chromatin organization, DNA repair, focal adhesion, and mitotic regulation. Enrichment of RAC3, RHOB, RHOC, and RHOF pathways suggests that Ref-1 inhibition may influence cytoskeletal organization, adhesion, and migratory signaling, while enrichment of nuclear speckles, RNA-splicing pathways, chromosome organization, and spindle-associated processes indicates that acute redox disruption rapidly engages RNA-processing machinery and nuclear stress responses before detectable changes in protein abundance (44–46). Because many kinases and phosphatases contain redox-sensitive cysteine residues, these phosphorylation changes likely represent downstream consequences of altered cellular redox homeostasis. This coordinated PTM remodeling was further illustrated by NF-κB1 and p53, two well-established Ref-1-regulated transcription factors (47). NF-κB1 exhibited oxidation at C61, C261, and C272 together with acetylation at K76, K78, K79, and K243 within or adjacent to the Rel homology domain. C61 is a well-characterized redox-sensitive residue that requires Ref-1-dependent reduction for efficient DNA binding, and its increased oxidation following APX2014 treatment is therefore consistent with inhibition of Ref-1 redox activity (28). Likewise, p53 displayed early oxidation of C275 and C277 within the DNA-binding domain followed by increased acetylation at K305, K357, K381, and K382 at later time points (48). The temporal separation between oxidation and acetylation in both proteins, together with the close spatial proximity of several modified residues, suggests that Ref-1 inhibition promotes coordinated PTM crosstalk rather than isolated modifications. Although the functional contribution of individual PTM sites remains to be determined, these findings demonstrate that Ref-1 inhibition rapidly remodels multiple regulatory layers on key transcription factors while simultaneously engaging broader signaling networks involved in stress adaptation and genome maintenance.

Unlike oxidation and phosphorylation, acetylation became substantially more prominent at the later time points, with enrichment of pathways involved in p53 regulation, histone acetylation, chromatin remodeling, DNA double-strand break repair, epigenetic regulation, and G2/M checkpoint control. This delayed response suggests that acetylation represents a downstream regulatory layer following the initial redox and phosphorylation changes, potentially reflecting altered chromatin regulation or changes in acetyltransferase/deacetylase activity and metabolic cofactors such as acetyl-CoA. Although the underlying mechanisms remain to be determined, the convergence of acetylation on chromatin-and DNA damage-associated pathways indicates that early redox perturbation is progressively translated into broader nuclear and transcriptional adaptation. These molecular changes were accompanied by functional consequences at later time points. Pathway enrichment identified chromosome segregation, mitotic nuclear division, and spindle-associated processes among the early PTM-responsive networks. These findings suggest that the early PTM remodeling observed after Ref-1 inhibition ultimately affects mitotic organization. Together, our findings support a temporally organized model in which Ref-1 inhibition rapidly induces widespread cysteine oxidation and phosphorylation before detectable changes in protein abundance occur, followed by progressive acetylation that reinforces chromatin remodeling and stress-response pathways. These coordinated PTM changes converge on mitochondrial metabolism, genome maintenance, and mitotic regulation, providing a mechanistic framework that links acute Ref-1 inhibition to the downstream metabolic and cellular phenotypes observed following prolonged treatment (**Fig. 7**).

**Figure 7.**
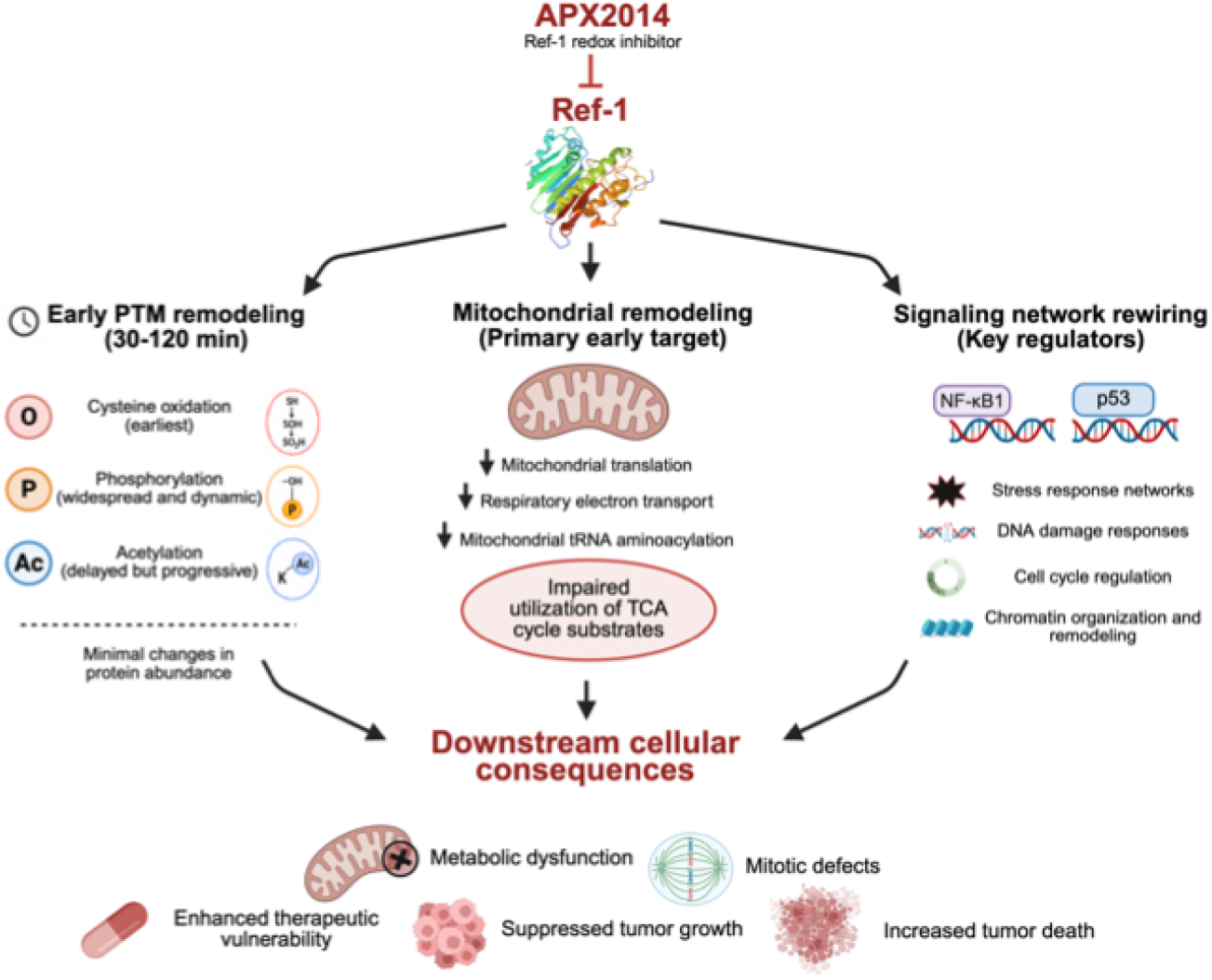
Summary model of the early cellular response to Ref-1 inhibition. Acute APX2014 treatment rapidly induces coordinated post-translational modification remodeling, including cysteine oxidation, phosphorylation, and acetylation, before detectable changes in protein abundance. Early PTM remodeling targets mitochondrial function and key signaling regulators, including NF-κB and p53, leading to altered stress-response pathways, impaired mitochondrial metabolism, mitotic defects, and downstream anti-tumor effects.

Although our findings provide a mechanistic framework for the early response to Ref-1 inhibition, additional studies will be needed to fully define the functional significance of these PTM changes. The present work captures the earliest molecular response to Ref-1 inhibition in a PDAC model, revealing a temporally organized sequence of redox, phosphorylation, and acetylation events that precedes changes in protein abundance. Determining whether this regulatory architecture is conserved across additional PDAC models and *in vivo* systems, and how individual PTMs contribute to mitochondrial dysfunction, transcriptional adaptation, and therapeutic response, will further refine our understanding of Ref-1 biology. Integrating these PTM datasets with complementary transcriptomic and metabolomic analyses especially under hypoxic conditions will also provide a more comprehensive view of how early signaling events evolve into the sustained phenotypic changes associated with Ref-1 inhibition.

## Conclusions

In conclusion, our integrated multi-PTM proteomic analysis demonstrates that pharmacologic Ref-1 inhibition initiates a rapid and temporally organized regulatory response that precedes detectable changes in protein abundance. Early cysteine oxidation and phosphorylation remodel mitochondrial, stress-response, and signaling networks, while later acetylation reinforces chromatin remodeling, p53 signaling, DNA-damage responses, and cell-cycle regulation. These coordinated molecular changes are followed by impaired mitochondrial substrate utilization and disruption of mitotic organization, establishing a mechanistic link between acute Ref-1 inhibition and the downstream metabolic and cellular phenotypes observed following prolonged treatment. Importantly, this study identifies mitochondria as a major early target of Ref-1 redox signaling and demonstrates that coordinated PTM remodeling of key regulatory proteins, including NF-κB1 and p53, represents an early mechanism by which Ref-1 inhibition disrupts tumor-promoting signaling networks. Together, these findings provide a systems-level framework for understanding the earliest cellular response to Ref-1 inhibition and further support the therapeutic potential of targeting Ref-1 in PDAC. These mechanistic insights also strengthen the translational rationale for the continued clinical development of Ref-1 redox inhibitors, building on the clinical experience with the first-in-class inhibitor APX3330 and informing the evaluation of next-generation compounds, including APX2014, as well as the development of PTM-based pharmacodynamic biomarkers and rational combination strategies.

## Declarations

### Ethics approval and consent to participate

Not applicable

### Consent for publication

Not applicable

### Availability of data and materials

All data generated or analyzed during this study are included in this published article. The mass spectrometry proteomics and PTM datasets (12 LC-MS runs for each datatype and 48.raw files in total) have been deposited in the ProteomeXchange (PXD082009) and can be accessed with the username MSV000102698 and the password PTM7243 (https://massive.ucsd.edu/ProteoSAFe/private-dataset.jsp?task=a61260cbabcd4ca8a29a5fa83030e54a).

### Authors’ contributions

Designed the study-S.G., X.L., W.J.Q., M.L.F., T.Z., and M.R.K.; Performed experiments, validation and analysis-S.G., X.L., J.B.T., M.A.G., R.K.C., W.J.Q., E.S.P., and T.Z.; Imaging and data analysis-E.S.P.; Writing-S.G., X.L., M.L.F., T.Z., and M.R.K.; Supervision and Funding Acquisition-M.L.F., W.J.Q., T.Z., and M.R.K. All authors reviewed the manuscript.

## Supporting information

Supp Data 1.Abundance_PTM_data

Supp Data 2. Significant Redox sites

Supp Data 3. Significant Phosphorylation sites

Supp Data 4. Significant Acetylation sites

Supp Data 5_Redox_Early_cluster 5

## Acknowledgements

Patricia S Miller at Pacific Northwest National Laboratory (PNNL) is thanked for assistance in fractionation of the peptide samples. Matthew E. Monroe (PNNL) is thanked for uploading the raw and processed proteomics datasets to ProteomeXchange.

## Supplementary Data

**Supplemental Data 1.** Global proteomics and PTMomics datasets in Pa03C cells

**Supplemental Data 2.** Significantly changed redox sites

**Supplemental Data 3.** Significantly changed phosphorylation sites

**Supplemental Data 4.** Significantly changed acetylation sites

**Supplemental Data 5.** Early redox responders to APX2014 in Pa03C cells

---

**Supplementary Figure 1.**
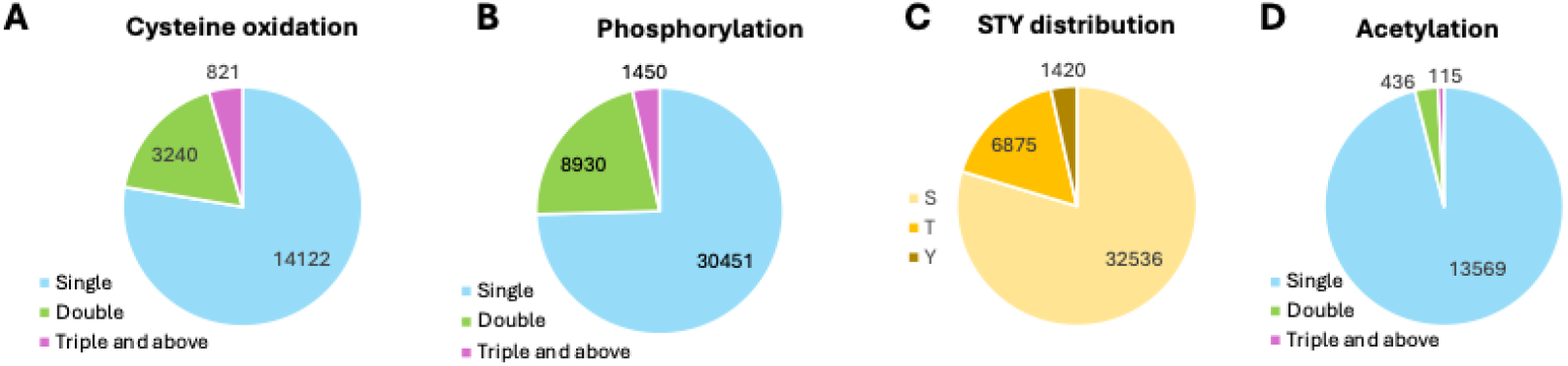
Distribution of PTM peptide classes quantified in the multi-PTM proteomics workflow. **(A)** Number of oxidized (redox) peptides containing single, double, or ≥3 oxidized cysteine sites; **(B)** Number of phosphopeptides containing single, double, or ≥3 phosphorylation sites; **(C)** Distribution of quantified phosphorylation sites by residue class (serine, threonine, tyrosine); **(D)** Number of acetylated peptides containing single, double, or ≥3 acetylation sites.

**Supplementary Figure 2.**
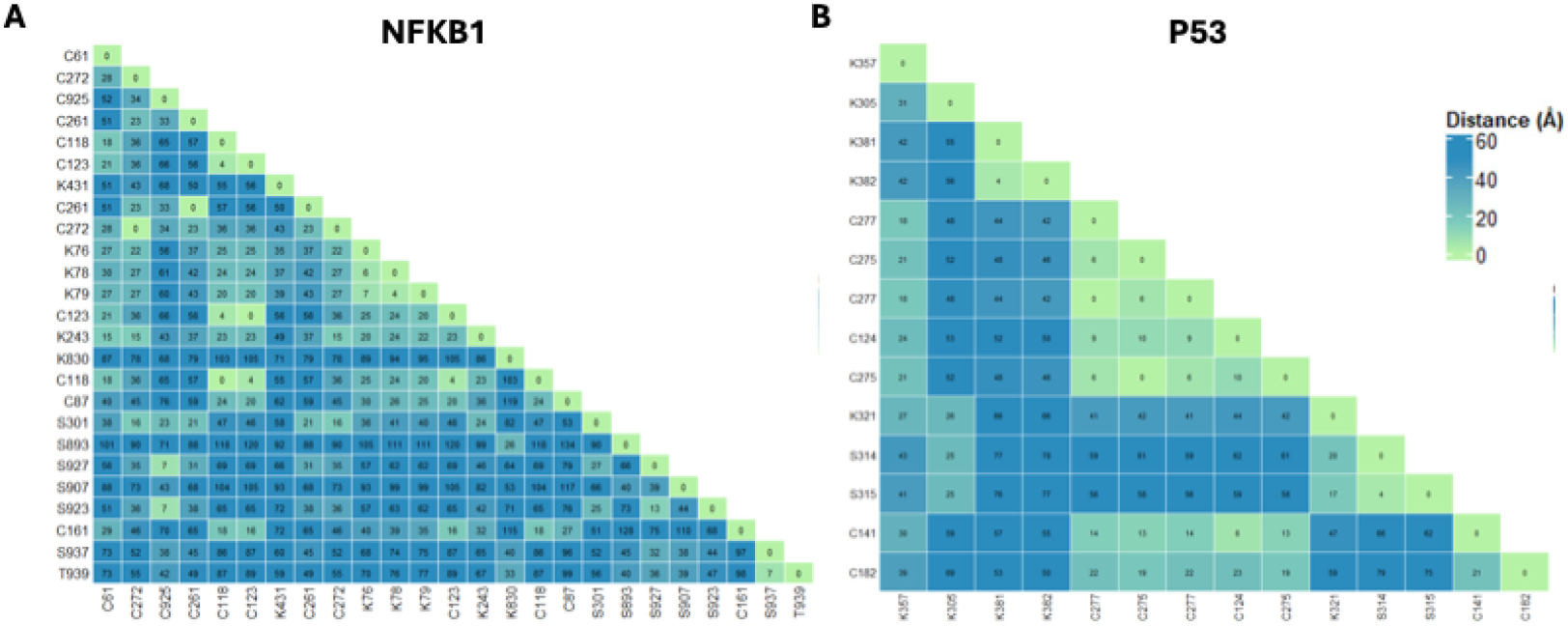
Spatial proximity of detected PTM sites on p53 and NF-κB1. Pairwise Cα– Cα distance matrix (Å) for quantified PTM sites mapped onto the AlphaFold structural model were provided for **(A)** NF-κB1 (NFKB1) and **(B)** p53 (TP53). Distances are color-coded as indicated, with smaller values reflecting spatial clustering of sites within the 3D structure.

**Supplementary Figure 3.**
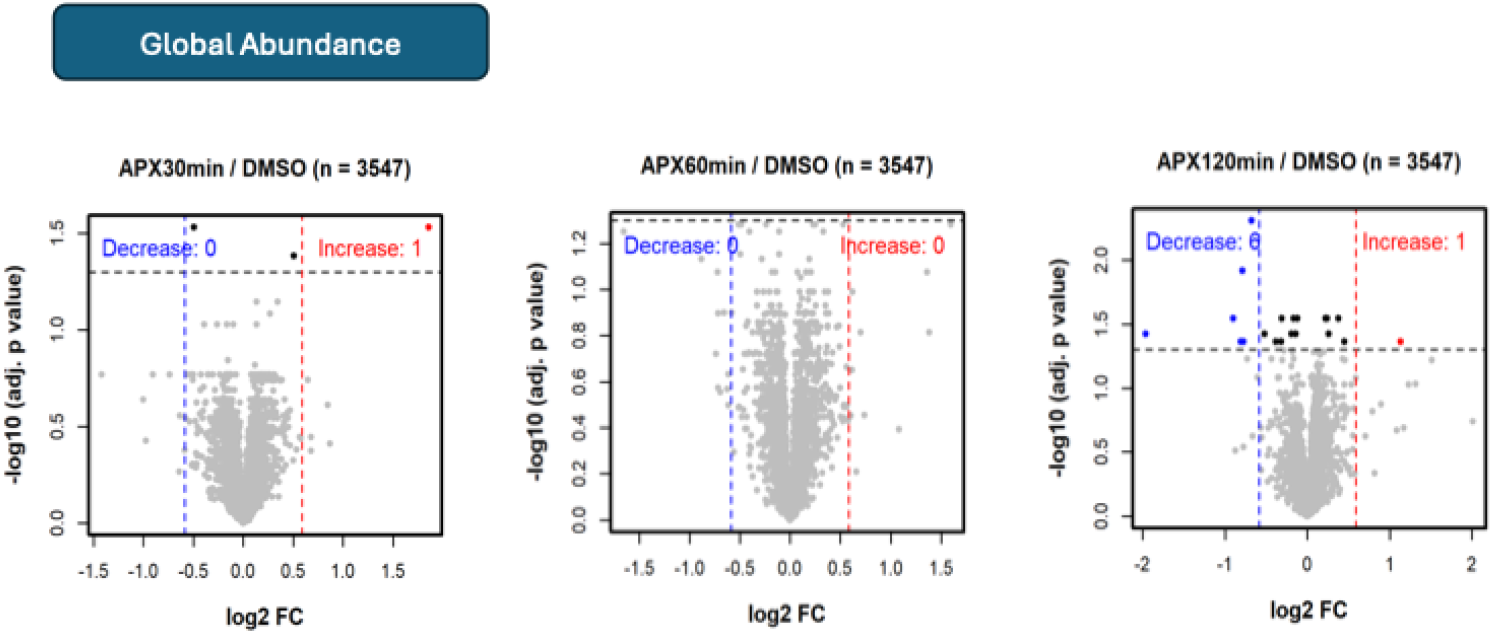
Differential proteome and PTMome analysis of APX2014 treatment. **(A)** Volcano plots showing global protein abundance changes (APX2014 vs DMSO) at 30, 60, and 120 min. Points are colored to indicate significantly decreased (blue) or increased (red) features, with non-significant features shown in gray/black; dashed lines denote fold-change and adjusted p-value thresholds. The number of quantified features (n) is indicated for each comparison.

**Supplementary Figure 4.**
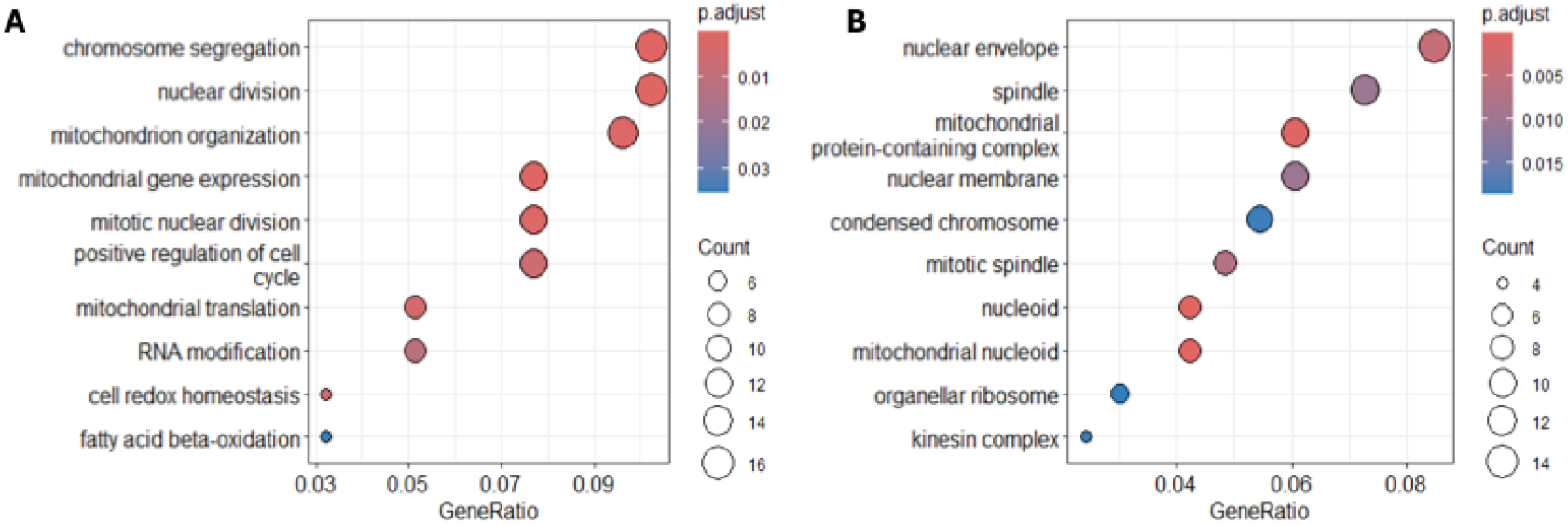
Functional enrichment of proteins exhibiting rapid redox-dependent changes after APX treatment. **(A)** Gene Ontology (GO) biological process enrichment shown as a dot plot; **(B)** GO cellular component enrichment.

**Supplementary Figure 5.**
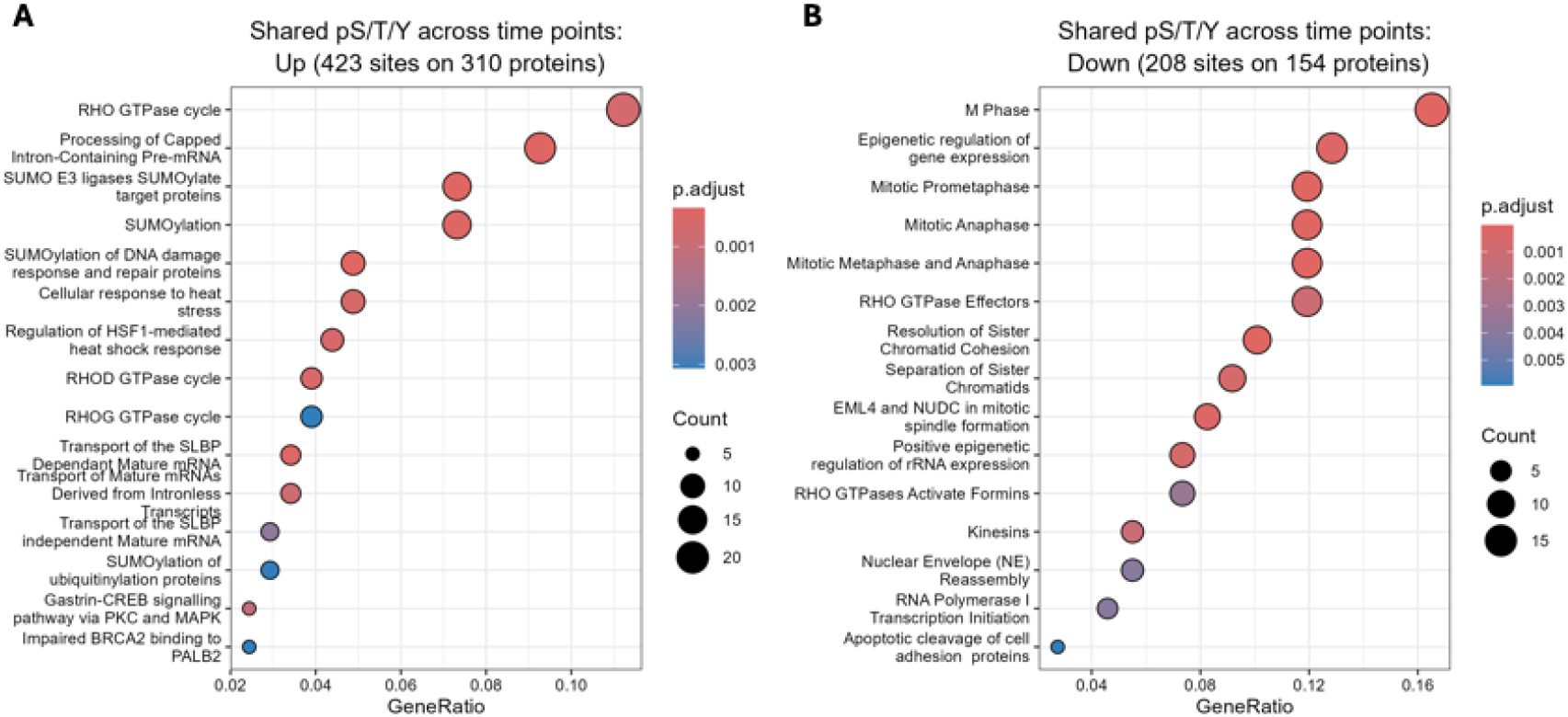
Reactome pathway enrichment of shared phosphorylation sites across time points. Reactome enrichment analysis was performed separately for phosphorylation sites that increased (**A**) or decreased (**B**) following APX2014 treatment. Dot size represents the number of proteins contributing to each pathway, the x-axis indicates the gene ratio, and color represents the Benjamini– Hochberg-adjusted *p* value. Pathways are shown among the 15 most significantly enriched terms for each group.

**Supplementary Figure 6.**
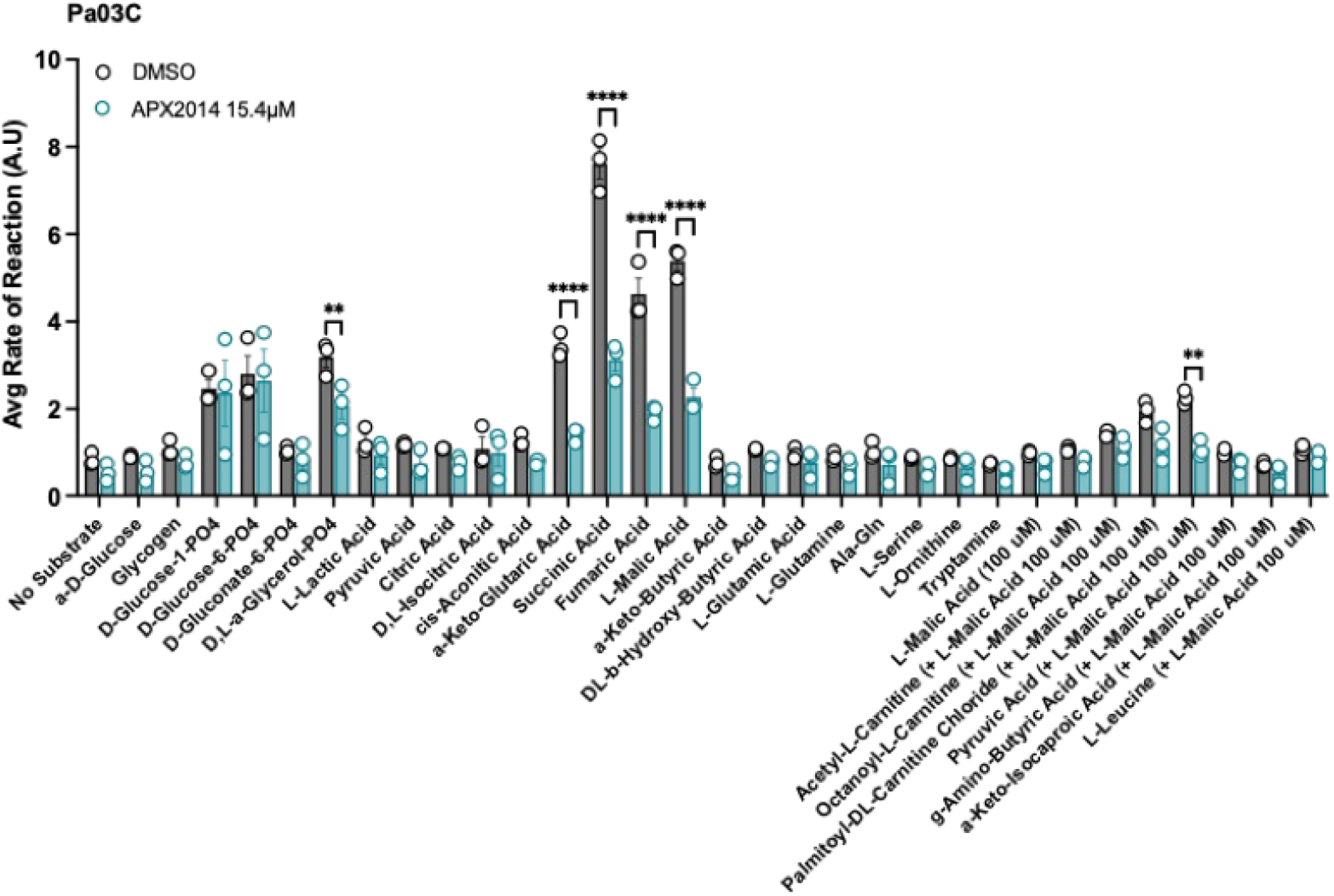
APX2014 selectively suppresses utilization of TCA cycle substrates in Pa03C pancreatic cancer cells. Pa03C cells were treated with DMSO or APX2014 (15.4 μM) for 24 h and average reaction rates (A.U) were calculated. Rates were calculated from the initial 90 min of the 240 min assay. The MitoPlate contains 31 mitochondrial substrates representing pathways involved in glycolysis, TCA cycle metabolism, amino acid metabolism, and fatty acid metabolism. Data are presented as mean ± SE from n = 3 independent biological replicates. Statistical significance was determined by two-way ANOVA. **p< 0.01, and ****p<0.0001.

